# Single Cell RNA-sequencing reveals cellular heterogeneity, stage transition and antigenic variation during stress adaptation in synchronized *Plasmodium falciparum*

**DOI:** 10.1101/752543

**Authors:** Mukul Rawat, Ashish Srivastava, Ishaan Gupta, Krishanpal Karmodiya

**Affiliations:** Department of Biology, Indian Institute of Science Education and Research, Dr. Homi Bhabha Road, Pashan, Pune 411 008, Maharashtra, India; Department of Biological Sciences, Indian Institute of Science Education and Research Bhopal, Indore By-pass Road Bhauri Bhopal, 462066, Madhya Pradesh, India

**Author notes:** Deceased on 06^th^ April, 2019 (Ashish Srivastava).

**Keywords:** Malaria, *Plasmodium falciparum*, Single cell RNA sequencing, Stress response, Antigenic variation, Transportation

## Abstract

The malaria parasite has a complex life cycle exhibiting phenotypic and morphogenic variations in two different hosts. Phenotypic cell-to-cell variability can be an important determinant of cellular adaptation, stress tolerance and immune evasion in the host. To investigate cellular heterogeneity, we performed single cell RNA-sequencing (scRNA-seq) of 4949 and 6873 synchronized *Plasmodium* cells in control and under temperature stress condition (phenocopying the cyclic bouts of fever experienced during malarial infection). High-resolution clustering of scRNA-seq datasets and a combination of gene signatures allow identification of cellular heterogeneity and stage transition during stress adaptation. We identified a subset of parasites primed for gametogenesis and another subset primed for stress adaptation. Interestingly, temperature stress inducted the process of gametogenesis by upregulation of master regulator (AP2-G) of sexual conversion. Moreover, pseudotime analysis indicated bifurcation for cell-fate decision to gametogenesis at two different stages of intra-erythrocytic cycle. Furthermore, we identified a rare population of cells, which is only emerged during the stress condition, showing the reactive state of the pathogen against the temperature stress condition. Interestingly, genes associated with the gametogenesis, chaperon activity and maintenance of cellular homeostasis showed maximum variation under temperature stress condition. We also developed an online exploratory tool (website: http://bit.ly/plasmo_sync), which will provide new insights into gene function under normal and physiological stress condition. Thus, our study suggests that the variability and versatility of the maintenance of cellular homeostasis should enable cells to survive under different stress conditions, and may act as an important stimulator of development of drug-resistance in *Plasmodium falciparum.*

## Introduction

Malaria is a major public health problem, with the parasite *Plasmodium falciparum* causing most of the malaria-associated mortality. Despite global efforts to eradicate the disease, over one million people die and 300-600 million suffer due to malaria every year [1]. *P. falciparum* completes its life cycle into two different hosts progressing through multiple developmental stages. The persistent infection and clinical manifestation of malaria require development of parasite in red blood cells (RBCs) via asexual intra-erythrocytic cycle (IEC). During the 48-h IEC, parasite invades RBC and develops into a ring stage, followed by trophozoite and schizont stages. The transmission of malaria parasite to mosquito depends on intra-erythrocytic differentiation to gametocytes in human host. Thus, parasite maintains balance in transmission and persistent infection by an unknown mechanism. Recent single-cell RNA sequencing (scRNA-seq) suggests commitment to gametocytes during the preceding cycles by epigenetic regulation of master regulator of sexual development, AP2-G [2-4]. However, the environmental cues and the hitherto mechanism therein, which initiate the gametogenesis are not yet clear.

*P. falciparum* is exposed to different environmental conditions during its life cycle in two different hosts. Moreover, it is also constantly exposed to different physiological conditions such as low glucose level [5] (hypoglycaemia), high temperature (fever) [6] and oxidative stress (haemoglobin degradation and drug therapy) [7] during IEC cycle. Under such adverse conditions some cells in the isogenic population survive whereas others do not. Presence of heterogeneity is indicated in both unicellular and multicellular organism at different levels and it helps in better survival of the organism during unfavourable conditions [8]. Multicellular organism are more advanced and developed in terms of fighting these fluctuations where heterogeneity is key to differentiation resulting in the diverse functions [9]. In unicellular organism presence of heterogeneity is a population level survival strategy [10]. Single cell organisms need to develop tight machinery for fighting against the unfavourable conditions. Slight change in the ability to respond to different stress conditions can result in competitive advantage over other organisms in the population. Ecological studies have suggested that survival chances of a particular population during extreme conditions are more if they have diverse populations [8]. Thus, cells which are less fit (stress-tolerant) in a particular population can have better survival chances during unfavourable conditions. But whether this stress-tolerant cells have a fully mounted stress response or they emerge from partial response is unclear. Thus, understanding cell-to-cell heterogeneity in stress survival has broad applications in stress adaptation and drug-resistance emergence.

One of the most common stresses experienced by *Plasmodium* is heat stress. During its intraerythrocytic life cycle, there are cyclic episodes of fever due to the release of merozoites. During the fever, temperature peaks as high as 41 degree Celsius for a period of 2-6 hours. Early ring stage of parasite growth is more resistant whereas later stages are more fragile to heat stress. High temperature during the malaria infection is known to be responsible for synchronizing *Plasmodium* growth in human host [11]. Despite of such variation in temperature during its life cycle, *Plasmodium* manages to overcome these unfavourable conditions resulting in successful infection. Stress response machinery is known to plays an important role in establishment of successful infection inside the host. Earlier, population studies have identified various proteins playing an important role in building temperature stress response [12]. Increase in temperature is also known to accelerate the growth and development of the parasites [6, 12]. Moreover, exposure of parasites to higher temperature results in increase of cytoadherence to CD36 and ICAM-1 receptors [12, 13]. This is attributed to the fact that during heat stress there is an increase in virulence (var) gene expression which helps in the binding of infected RBCs to the endothelial receptors resulting in cytoadherence [12]. Recent reports also suggest stress responses as crucial mediator of artemisinin resistance in *P. falciparum* [14, 15].

*Plasmodium* is known to have tight gene regulation orchestrated at multiple levels by chromatin structure [16, 17], transcription [18-20] and translation [21, 22], which helps in timely expression of different genes required at various stages of parasite growth [23]. Despite tight-gene regulation there is a huge amount of transcriptional heterogeneity exists in the parasite populations [3]. The cell-to-cell heterogeneity can be attributed to various environmental and physiological conditions such as availability of nutrients, temperature fluctuations, cell-cycle progression and different morphological states. Advent of single-cell RNA sequencing (scRNA-seq) has been instrumental in unravelling the cell-to-cell heterogeneity in the population [24-26]. It provides information pertinent to different states co-exist in a cell population leading to dynamic cell-to-cell variability in mRNA numbers and concentrations. Earlier study in *Plasmodium* using scRNA-seq has revealed the heterogeneity in the parasite population during intraerythrocytic like cycle [3].

In this study, we have performed scRNA-seq under control and temperature stress condition to understand the cell-to-cell heterogeneity within and between the *Plasmodium* populations. Using this approach, we identified a combination of gene signatures of cellular heterogeneity and stage transition during stress adaptation. Interestingly, we identified a subset of parasites primed for gametogenesis and another subset primed for stress adaptation. On temperature stress there is upregulation in the expression of gametocyte related makers in these parasites which indicates that high temperature can further trigger/accelerate the process of gametogenesis. Moreover, we identified a rare population of cells, which is only emerged during the stress condition, showing the reactive state of the pathogen against the temperature stress condition. Thus, our scRNA-seq analysis reveals important insights into the cell-to-cell heterogeneity in parasite population under temperature stress condition that will be instrumental towards mechanistic understanding of cellular adaptation, population dynamics and drug-resistance emergence in *P. falciparum.*

## Results

### Single-cell RNA sequencing (scRNA-Seq) during temperature stress in *P. falciparum*

*Plasmodium* while harboured within the human RBCs, exposed to different kinds of environmental and physiological stresses. The impact of these environmental and physiological changes on gene expression heterogeneity is important to understand the mechanism of stress tolerance and adaptation. To investigate this, we performed single-cell RNA sequencing (scRNA-seq) under control and physiologically stress condition, high temperature (equivalent to fever) in *P. falciparum*. Parasites were synchronized by sorbitol during early (0-3 hrs) ring stage. Temperature stress was given at 40 degree Celsius approximately at late ring stage (∼17 hrs) for a period of 6 hours. For parasites grown under normal and temperature stress conditions, the infected RBCs were enriched by Percoll gradient in order to get rid of uninfected RBCs. We obtained around 80% enrichment of infected RBCs using Percoll gradient centrifugation. Enriched parasites under normal and high temperature growth conditions were assayed for their single cell gene expression profiles using the 10x chromium single cell RNA-sequencing v3 chemistry (Figure 1A). We sequenced 4949 single cells in control and 6873 single cells under temperature stress condition. Median number of genes detected per cells (count ≥1) is 637 and 546 with sequencing depth (mean reads per cell per gene) 17945 and 13001 for control and temperature stress sample, respectively (Figure 1B).

**Figure 1:**
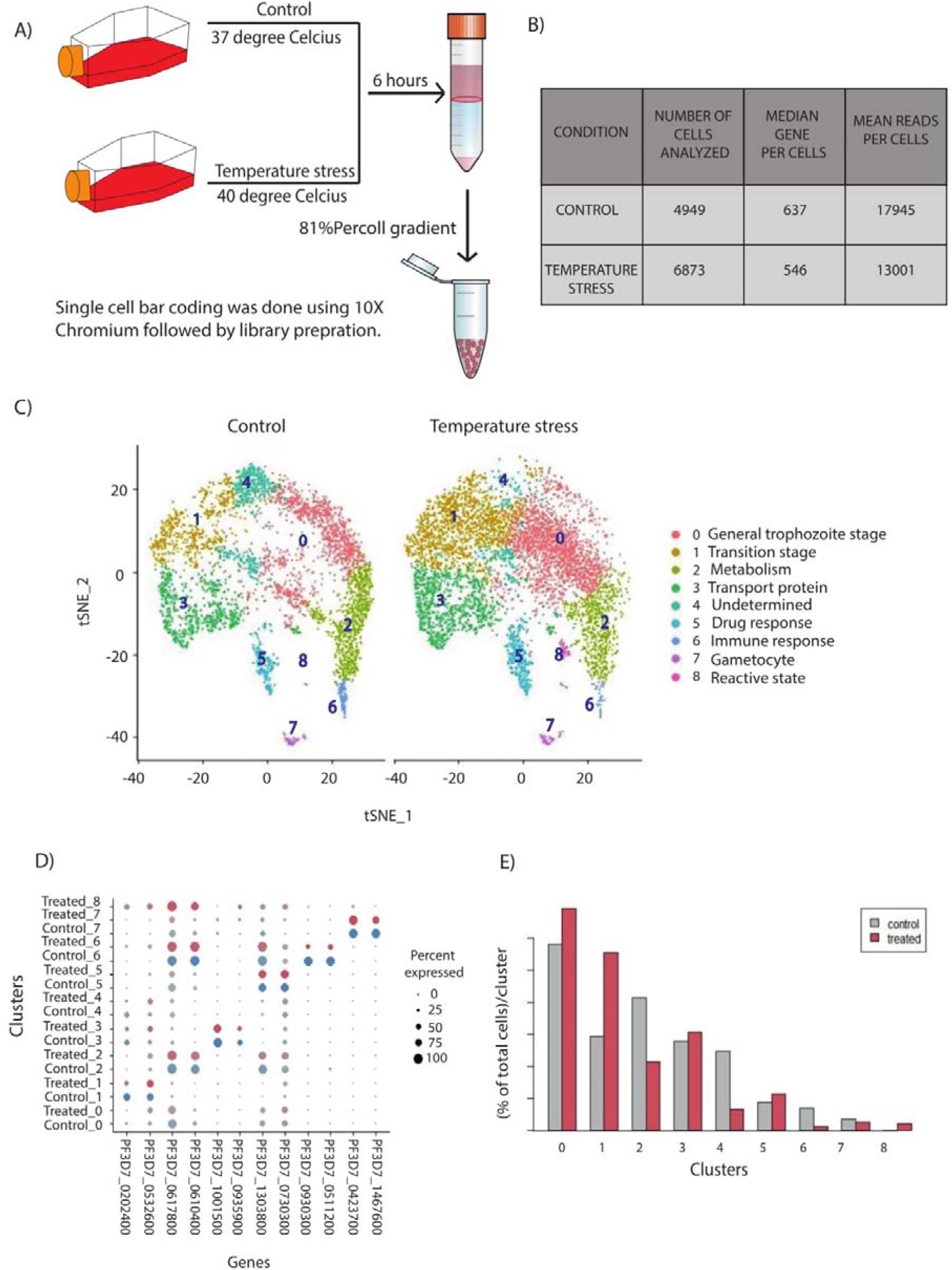
Single-cell RNA sequencing (scRNA-Seq) during temperature stress in *P. falciparum*. (A) Schematic representing the pipeline used for stress induction, isolation of single parasites and library preparation for scRNA-seq. Infected RBCs were enriched using 81% percoll gradient centrifugation. Single cell bar coding and library preparation was performed using 10X chromium protocol as per the manufacturer’s instruction. (B) Table stating the number of cells sequenced, median genes per cell and mean reads identified per cell under both normal and temperature stress conditions. (C) t-SNE plot representing the different clusters generated for the *Plasmodium* population in control and temperature stress condition. (D) Bubble plot showing the percentage of cells expressing a marker gene under control and temperature stress condition. Marker genes are specific to each cluster and expressed by at least 30% of the cells and have 50 % or higher average expression than other clusters. (E) Number of cells in different clusters during control and temperature stress condition. Number of cells increased in cluster 0, 1, 3 and 5 and decreased in cluster 2, 4, 6, 7 and 8.

Further to identify different cell types (Figure 1C), markers genes specific to each cluster identified, which are expressed by at least 30% of the population and have 50% higher average expression than other clusters (Figure 1D, Supplementary Figure S1A-S1B and Supplementary table S1). Interestingly, we found a novel cluster which is only present during the temperature stress conditions (Figure 1C). Moreover, we also found significant change in the number of cells (%) in different cluster during temperature stress condition (Figure 1E). Changes in the number of cells maybe indicative of how the treatment rewired gene expression and biases the cells to exist in specific states that may be functionally relevant to the treatment (particularly evident for cluster 0, 1 and 4). Interestingly despite of double synchronization of the parasites we were able to observe significant variation in the population of the parasites during control condition. This is an important observation since most of the experiments performed in the field depend on the usual method of synchronization like percoll gradient and sorbitol treatment. In summary, scRNA-seq datasets generated under control and temperature stress conditions indicative of transcriptional heterogeneity in *P. falciparum*, which is analysed in detail below.

### scRNA-seq identifies transcriptome heterogeneity during temperature stress condition

To understand the cell-to-cell variability of transcriptome regulation we calculated average number of RNA molecule per cell during control and temperature stress condition. Interestingly, average numbers of RNA molecules per cells are downregulated during the temperature stress condition indicative of the fact that parasites slowdown or shutoff global transcription machinery during temperature stress condition (Figure 2A). Downregulation can be also observed when RNA molecules per cells are calculated for each cluster except cluster 1 (Figure 2B). The number of RNA molecules is particularly affected in clusters associated with stress tolerance, immune response, and gametogenesis (cluster 5-7). Next, we performed bulk RNA sequencing under control and temperature stress conditions to compare global transcription. We observed high correlation in relative expression in control (Spearman’s correlation, 0.51) and during temperature stress condition (Spearman’s correlation, 0.41) in scRNA-seq and bulk RNA sequencing. Moreover, we also observed global transcriptional downregulation during temperature stress condition in bulk RNA-sequencing (Figure 2C). Further to understand the heterogeneity present between the control and temperature stress condition, we calculated the measure of dispersion with in a population using Coefficient of Variation (CV). To check the heterogeneity between the populations we performed the Flinger-Killeen test which checks for homogeneity for variance when data is non-normally distributed. The Flinger-Killeen Test [27] for equal coefficients of variation suggest significant variation between control and stress condition (CV, 128.19 and 136.32 for control and temperature stress condition, respectively; p value =2.08e-5) (Figure 2D). Interestingly, genes associated with the gametogenesis, chaperon activity and maintenance of cellular homeostasis showed maximum variation under temperature stress condition (Figure 2E and 2F). This in turn suggests that variability and versatility of the maintenance of cellular homeostasis should enable cells to survive under different stress conditions, and may act as an important stimulator of development of drug-resistance in *P. falciparum*.

**Figure 2:**
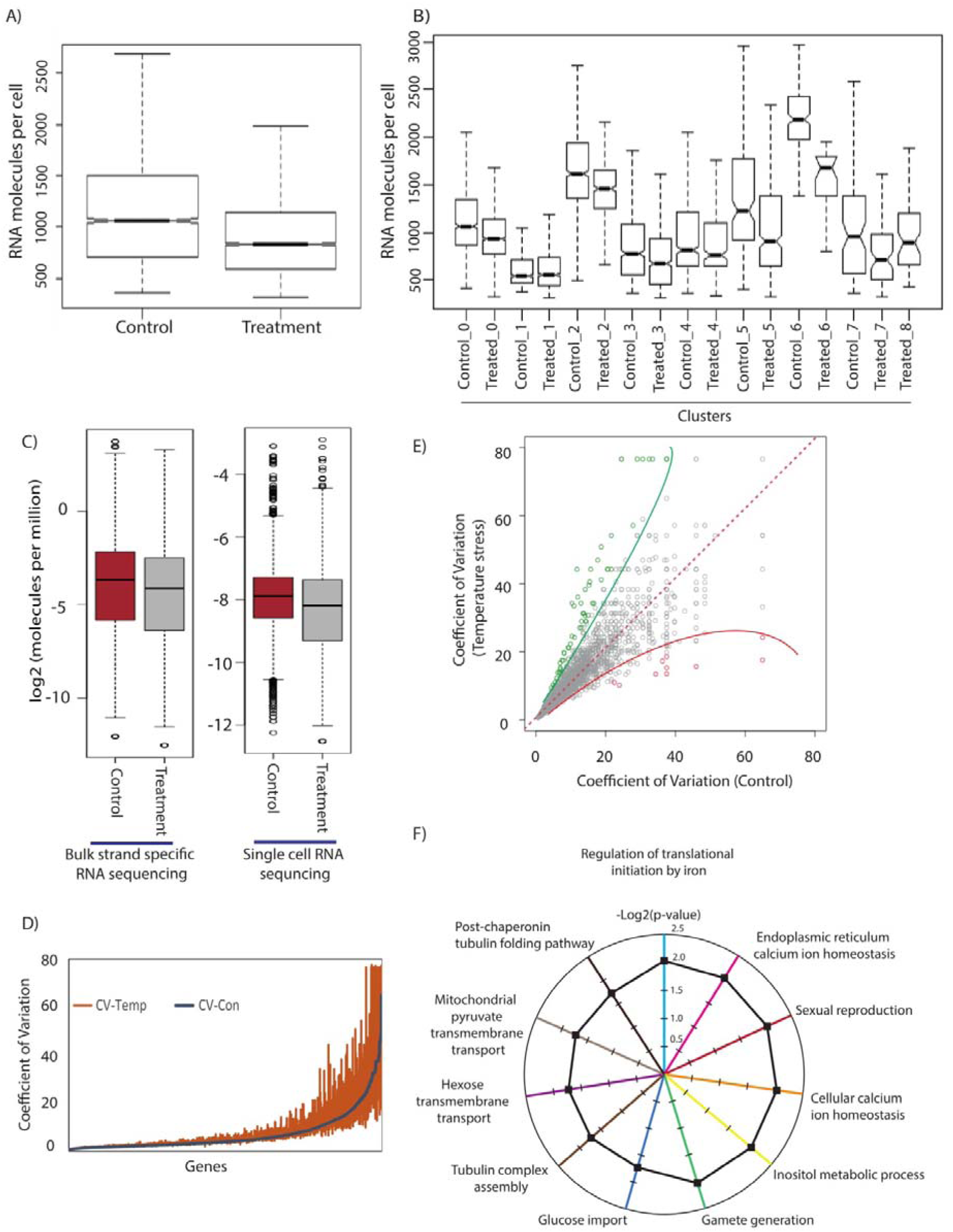
scRNA-seq identifies transcriptome heterogeneity during temperature stress condition. (A) Average RNA molecules expressed per cell is plotted for cells under control and temperature stress condition. Downregulation in the number of RNA molecules was observed during temperature stress. (B) Average number of RNA molecules were plotted for each clusters. (C) Mean expression level of all the genes expressed during control and temperature stress condition in both single-cell RNA sequencing and bulk-RNA sequencing. Overall transcription is suppressed upon temperature stress condition. (D, E) The Flinger-Killeen Test for equal coefficients of variation suggest significant variation between control and temperature stress condition (CV, 128.19 and 136.32 for control and temperature stress condition, respectively; p value =2.08e-5). Coefficients of variation are plotted for temperature stress condition upon control condition. Under temperature stress condition genes associated with cellular homeostasis show greater variation. (F) Radar plot showing the gene ontology terms enriched for the genes which show highest variation under stress conditions.

### Gene expression analysis of stress responsive genes under temperature stress condition

Most of the organisms have evolutionary conserved mechanism of stress response against variety of the environmental fluctuations. These stress response machineries consist of various proteins maintaining cellular homeostasis during unfavourable conditions. One of the most studied classes of proteins which play a crucial role in maintenance of cellular homeostasis is heat shock proteins. These proteins are known to play diverse functions like refolding of misfolded proteins and helps in the mRNA processing and maturation. They are considered central to the stress response machinery in higher eukaryotic systems. In order to understand the transcriptional variation in stress responsive genes under temperature stress condition we plotted expression levels of stress responsive genes relative to ribosomal protein genes (which were found to be unchanged in all clusters). Though most of the clusters showed upregulation in the expression of stress responsive genes, clusters associated with stress and immune response, and gametogenesis (cluster 5-7) showed highest overall expression and upregulation under temperature stress condition (Figure 3A and 3B). Interestingly, cluster 8, which is unique to temperature stress condition, showed very high expression of antigenic variation gene-families like var, rifins and stevors. This unique cluster represents cells which express the genes associated with the reactive state of the pathogen against the elevated temperature. Furthermore, we looked at the different stage specific markers for these clusters. Cluster 3 showed the genes specific to ring stage markers whereas cluster 6 shows genes specific to schizont stage (Figure 3C). Marker genes from cluster 6 shows expression of different chaperone proteins which are known to play an important role in the stress response in an organism (Figure 3C). Gene ontology of the marker genes from these clusters indicate that they play an important role in stress response and immune response functions. Most of the other cluster shows gene expression similar to trophozoite stage which is expected since parasites were harvested during trophozoite stage. This indicates that the heterogeneity is an inherent property of *P. falciparum* growth as it is also exhibited under control condition. This is interesting since we have used synchronized parasites for the study. In order to further validate our observation we compared our clustering with the single cell data available from Malaria Cell Atlas [28]. Since the Malaria Cell Atlas has asynchronized parasites we observed a partial overlap with their data (Supplementary Figure S1C). Around 98% of cells of our datasets (both control and temperature stress condition) cluster together with cells of data from Malaria Cell Atlas that lie in clusters containing at least 0.5% cells from the our datasets on the other hand only 49.8 % of the cells of the Malaria Cell Atlas dataset cluster together with cells of our datasets that lie in clusters containing at least 0.5% cells from our dataset.

**Figure 3:**
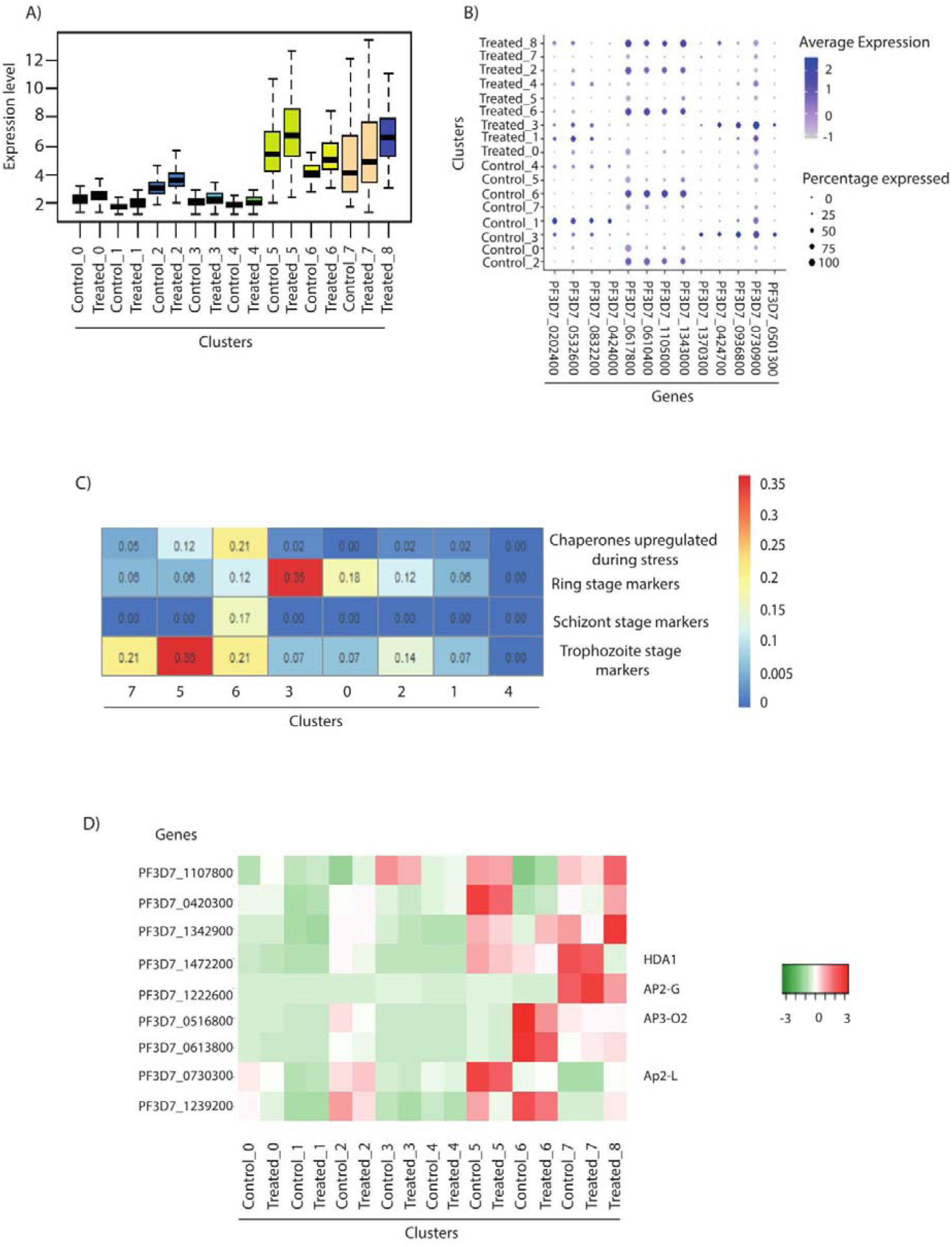
Gene expression analysis of stress responsive genes under temperature stress condition. (A) Change in the expression level of the stress responsive genes in different clusters. Gene expression is plotted over ribosomal proteins since expression of these genes was found to be relatively uniform in all the clusters. (B) Bubble plot showing the percentage of cell and average expression of a particular gene in different cluster. Stress responsive genes were plotted to show gene expression in different clusters. (C) Heatmap showing the expression of stage specific gene markers in different clusters. Different stages of parasite growth can be identified using these stage specific markers in various clusters. (D) Heat map showing the specific AP2 transcription factors are enriched in different clusters indicating that these transcription factors may be regulating the expression of cluster specific gene expression programs.

We wondered if there is any specific transcription factor associated with the regulation of stress and immune response clusters (5-7). We looked at the status of AP2 transcription factors in these different clusters (Figure 3D). Interestingly, we identified an AP2 transcription factors (PF3D7_1239200), which is downregulated in cluster 5. Another AP2 transcription factor (PF3D7_1342900) was specifically found to be upregulated in cluster 6 during the temperature stress condition. PF3D7_1342900, AP2 transcription factor was also found to be highly expressed in other stress responsive clusters like 7 and 8 (Figure 3D). Other AP2 transcription factors with cluster specific expressions are represented in the heat map (Figure 3D). Thus, it is plausible that these marker AP2 transcription factors regulating the expression of the genes which are differentially expression in their respective clusters. Interestingly we identified histone deacetylase, HDA1 (PF3D7_1472200) showing high expression in cluster 5, 6, 7 (Figure 3D). Expression of the histone deacetylase, HDA1 decreases upon temperature treatment suggestive of the fact that it may help in the upregulation of genes related to stress response and gametogenesis

### Temperature stress results in induction of gametogenesis in *P. falciparum*

Gametogenesis is a genetically encoded phenomenon but it is known to be influenced and triggered by various environmental fluctuations [27]. Previous studies have suggested various factors such as host immune response, high parasitemia load, elevated temperature, drug treatment etc., act as trigger for the process of gametogenesis in *P. falciparum* [29-34]. Furthermore, stress conditions are also shown to induce gametogenesis in *Plasmodium* [35]. In similar context, we investigated the enrichment of gametocyte markers in various clusters under temperature stress condition (Figure 4A). Interestingly, we found significant upregulation of gametocyte marker (PF3D7_1302100, PF3D7_0406200, PF3D7_1253000) as well as gametogenesis regulator (PF3D7_1222600) genes in cluster 7 (Figure 4B). Interestingly, genes which are only expressed in cluster 7 include PF3D7_1222600 (AP2 domain transcription factor AP2-G), PF3D7_1466200 (early gametocyte enriched phosphoprotein EGXP), PF3D7_0936600 (gametocyte exported protein 5), PF3D7_1253000 (gametocyte erythrocyte cytosolic protein), PF3D7_1302100 (gamete antigen 27/25) and PF3D7_0406200 (sexual stage-specific protein precursor) (Figure 4C). Recent report in *P. falciparum* identified markers of gametogenesis using scRNA-seq data [2]. We observed significant overlap of these genes, which are upregulated in the gametocyte specific population like surface-associated interspersed protein 8.2 (SURFIN 8.2) (PF3D7_0830800), early transcribed membrane protein 4, serine/threonine protein kinase (PF3D7_1356800), putative NYN domain-containing protein (PF3D7_0406500), putative CPSF (cleavage and polyadenylation specific factor) (PF3D7_0317700), PF3D7_0801900 (lysine-specific histone demethylase) and PF3D7_1472200 in cluster 7 (Supplementary Figure S1A and Supplementary table S1). Thus, we hypothesize that cluster 7 represent a rare population within the parasites, which is already primed for gametogenesis and these parasites show induction of gametogenesis once they are exposed to the higher temperature.

**Figure 4:**
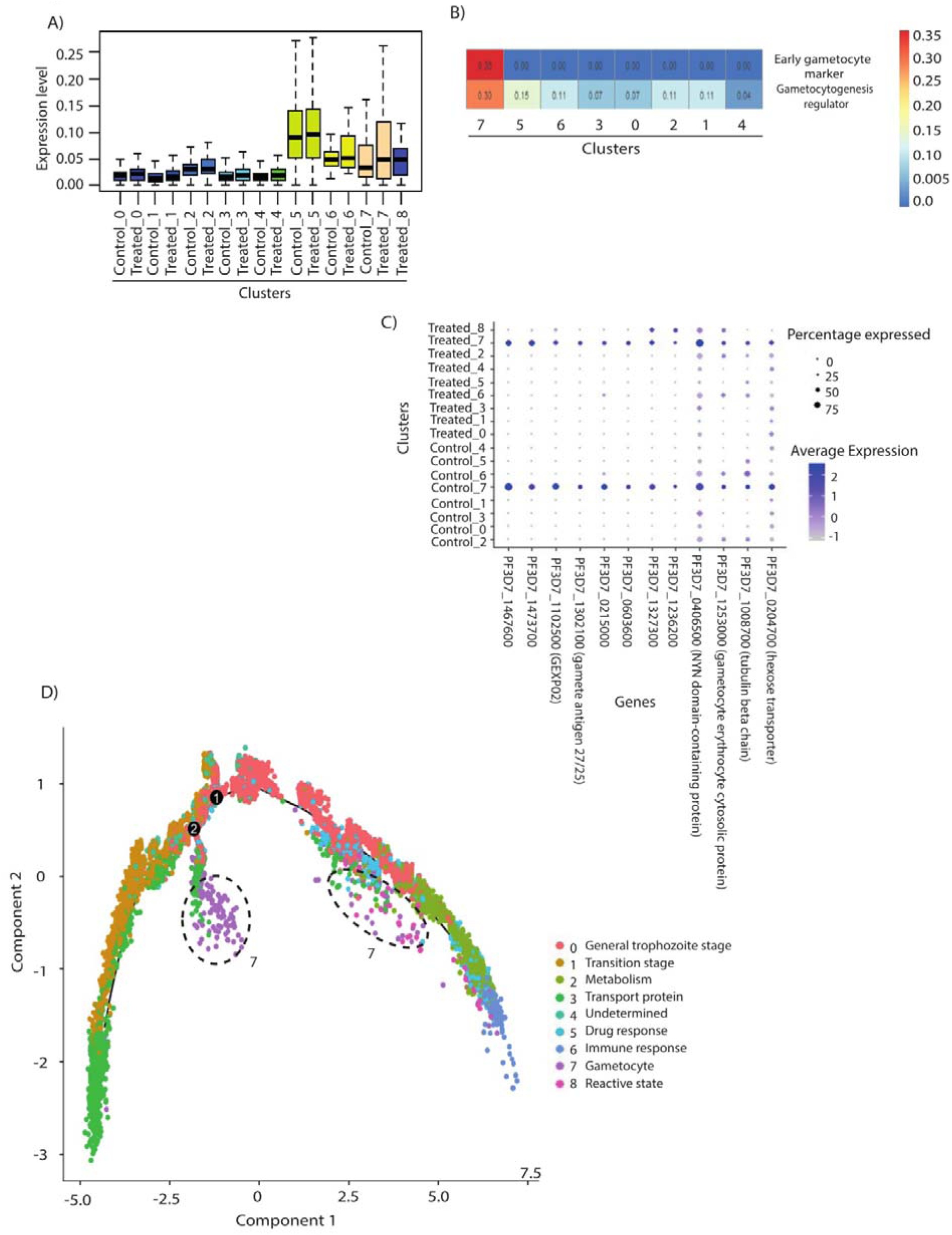
Temperature stress results in induction of gametogenesis in *P. falciparum*. (A) Change in the expression level of the gametogenesis related genes in different clusters. Gene expression is plotted over ribosomal proteins since expression of these genes was found to be relatively uniform in all the clusters. (B) Heat map showing the expression of early gametocyte marker and gametocyte regulator in different clusters. Stage specific gene markers were used to identify the stage of the parasites in different clusters. (C) Bubble plot showing the percentage of cell and average expression of gametocyte related genes. (D) Pseudotime analysis of the parasites sequenced under control and temperature stress condition. Pseudotime analysis indicated bifurcation at two stages of parasite growth for gametogenesis.

Furthermore, we performed pseudotime analysis across the intraerythrocytic life cycle of *P. falciparum* to understand the cell-fate decisions and stage-transition under temperature stress condition (Figure 4D). Pseudotime ordering reveals a gradual stage-transition from cluster 3 (early ring stage) to cluster 6 (trophozoite/early schizont) in control as well as temperature stress condition. Interestingly, we observed two branching (indicated by 1 and 2 in Figure 4D). Branch 1 indicates parasites which failed to diverge successfully from the default development trajectory. While second branch point indicates the parasites in cluster 7 which are primed for gametogenesis (expressing markers of gametogenesis) under control and temperature stress condition. Moreover, there is significant upregulation of gametogenesis related genes upon temperature stress condition (Cluster 7 in Figure 4A). This, in turn suggests that, though parasites are primed for gametogenesis even in the control samples, temperature stress trigger the process of gametogenesis (Figure 4A). Few parasites primed for gametogenesis are also found in the trophozoite stage of parasite development. Thus, it is possible that a subset of parasites is primed for gametogenesis and cell-fate decisions to gametogenesis and this process can be triggered depending on the environmental conditions.

### Export proteins regulation plays an important role in stress response adaptation

*Plasmodium* during intraerythrocytic life cycle exports various proteins on the surface of the infected erythrocyte. Many of the exported proteins belong to the multivariant gene family, helping the parasites in sequestration by binding to various endothelial receptors. Also, there are export proteins mediate the process of rosetting, which helps in avoiding exposure of the parasite to host immune response. Previous studies on gene expression using microarray have suggested that there is a huge increase in the proteins which are known to be either transmembrane or secreted proteins during the temperature stress in *P. falciparum* [12]. Most of these proteins have either PEXEL motif or host target signal sequence. Such transportation is usually mediated during the early stages of intraerythrocytic life cycle [12]. Though transported proteins expressed in almost all clusters, they are significantly upregulated in cluster 3 (Figure 5A and 5B). This is intuitive as most of the transportation related processes take place during ring stage (and cluster 3 also exhibits characteristics of ring stage parasites, Figure 3C). Some of the export proteins which are upregulated in cluster 3 include PfEMP1 trafficking protein (which helps in the trafficking of PfEMP1 protein on RBC surface) and PF3D7_0424700 (FIKK kinase), which mediate the virulence associated changes over *Plasmodium* infected RBC surface. Interestingly, cluster 3 also shows downregulation of some of the transport proteins like PF3D7_1370300, PF3D7_0730800 and PF3D7_0532600 etc. This indicates mutually exclusive preference for transportation of proteins during the temperature stress conditions (Figure 5C and 5D). Various transport proteins showing deregulation during the temperature stress are represented in the heat map plotted for each clusters (Figure 5E). Surprisingly, cluster 1 also shows overrepresentation of genes which helps in transportation of proteins in parasites (Figure 5B). This might be the transition cluster, originated from cluster 3 since parasite of these stage show expression of both ring and trophozoite stage gene expression and helps in transportation of different proteins. Presence of the cluster 1 adjacent to cluster 3 in pseudotime analysis (Figure 4D) further supports our observation that cluster 1 and cluster 3 parasites show similarity in their gene expression profile.

**Figure 5:**
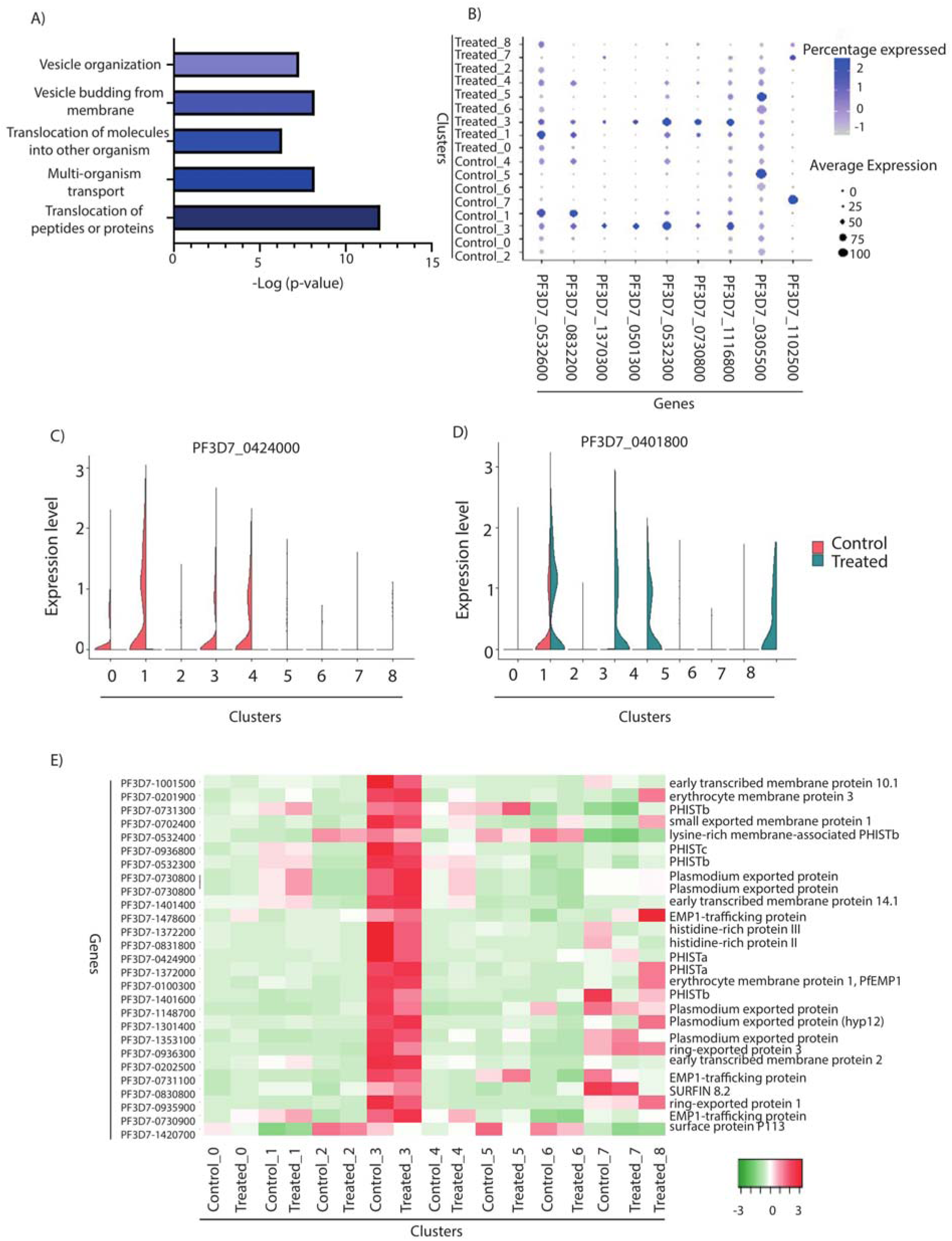
Export proteins regulation plays an important role in stress response adaptation. (A) Gene ontology analysis of the markers genes of cluster 3 indicated that these parasites have higher expression of genes related to transportation and secretion. (B) Bubble plot showing the percentage of cell and average expression of transported proteins. (C, D) Transport related proteins showing deregulation during temperature stress condition. Mutually exclusive expression of representative transport proteins in control and temperature stress condition. (E) Heat map showing the change in the various transport proteins during the temperature stress in different clusters.

### Virulence gene regulation during temperature stress condition

*Plasmodium falciparum* encodes several clonally variant multicopy gene families such as var, rifin and stevor, which are upon expression, presented on the surface of infected RBCs. These proteins play a central role in enabling host immune evasion and promoting pathogenesis. PfEMP1 (*Plasmodium falciparum* Erythrocyte membrane protein 1) is the most commonly expressed immunodominant antigen on iRBCs. PfEMP1, encoded by a family of 60 var genes, believed to exhibit antigenic switching upon immune exposure and/or environmental fluctuation, stress conditions or chromatin organization etc. [36-39]. Bulk RNA-sequencing studies have suggested expression of only one var gene in a population. However, the mechanism of selection of a single var gene for expression and suppression of rest of the var genes is still not clear. Recent study using scRNA-seq employing Smart-seq2 has looked into the expression of var genes at the ring stage of the parasite [3]. As per this study majority of parasites (13 out of 17 individual cells) show expression of one dominant var gene belonging to var sub group B. Furthermore, only a few parasites showed expression of two var genes per cells [3].

We decided to look at the expression of var genes per cell using the scRNA sequencing data under stress condition (Figure 6A and 6B). Interestingly, we found that var genes are expressed in cluster 3 (transport related cluster) and in cluster 8 (reactive stage of pathogen to elevated temperature) (Figure 6A). We further plotted the normalized expression level of different var genes in control and temperature conditions and found the few var genes are expressed at a higher level in control condition. On temperature treatment, more var genes are expressed at higher expression level than control conditions (Figure 6B). However, it is not clear if multiple var genes are expressed per cell or per parasite population. To investigate the regulation of expression of var genes at single cell level, we performed targeted analysis of var gene expression under control and temperature stress condition. We observed 1-4 var gene expression per cell under control condition and it goes up to 1-6 var gene expression per cell under temperature stress condition (Figure 6C). It is possible that parasites in order to survive the unfavourable conditions may express more than one var genes per cell to increase the chances of binding to the endothelial cells. Interestingly, basal level of transcription was high for many var genes under temperature stress condition whereas it was comparable to control condition at higher expression level (Figure 6D). Thus, it can only be speculated that parasite under unfavourable conditions express multiple var genes at basal levels and only a few of them are selected for high expression. Future functional analyses will discern weather var gene selection and activation involves poised chromatin activation or var-gene coded intronic non-coding RNAs. We next investigated the expression levels of var genes in scRNA-seq data from control and temperature stress condition. Interestingly, we observed that var group B and C are highly expressed under both the conditions. Moreover, bulk RNA-sequencing also showed upregulation of var genes during temperature stress condition which are also validated RT-qPCR (Figure 6E). Expression level of var genes was quantified after 6 hours heat treatment. Expression level was also checked after 54 hours to check the effect of heat treatment at later hours. Interestingly, we found var group B gene Pf3D7_1200100 to be highly upregulated in control sample. We wondered if expression of var group B is higher in patient samples as indicated by this and earlier study [3] and if it is geographical location dependant. Interestingly, we observed that indeed var group B is highly expressed in patient samples and it is independent of geographical location as we have observed similar profile from African and South Asian patient samples (Supplementary Figure 2A – 2C). This indicates that there is preference for expression of var group B over other var groups in *P. falciparum*. This can be further exploited to specifically design vaccines targeting members of var group B in *P. falciparum*.

**Figure 6:**
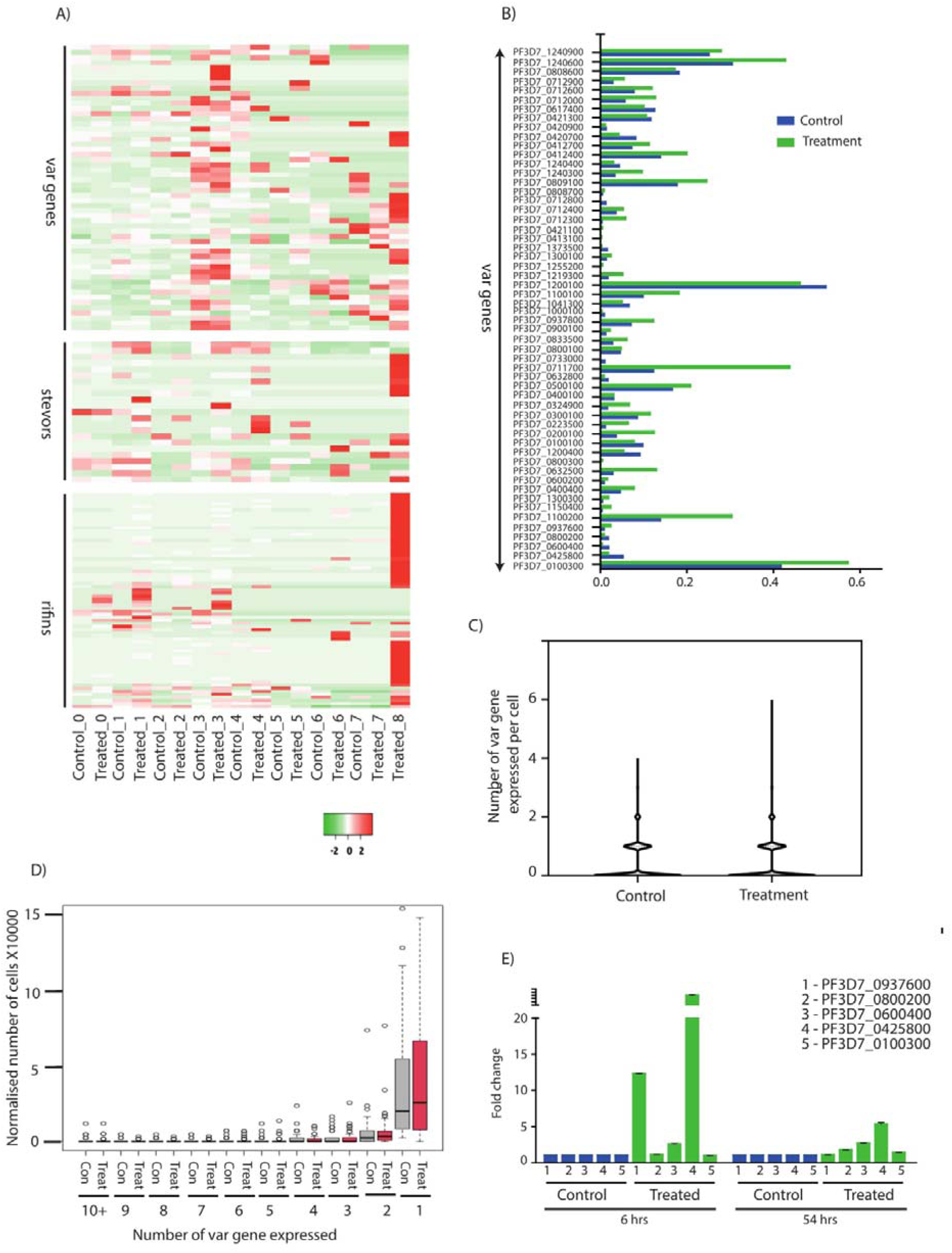
Virulence gene regulation during temperature stress condition. (A) Heat map showing the normalised expression of various var, rifins and stevors genes during the temperature treatment in different clusters. (B) Bar plot showing the upregulation of different var genes during temperature treatment. (C) Violin plot indicating the number of var genes expressed per cell. During control cells show 0-4 var genes per cell which is increased to 0-6 var genes per cells under temperature stress condition. (D) Expression level of var genes in control and temperature stress condition. Basal level of transcription was high for many var genes under temperature stress conditions whereas it was comparable to control condition at higher expression level. (E) RT-qPCR for parasites under temperature treatment for var genes indicating the upregulation of var genes during temperature stress condition. Heat stress was given for 6 hours and expression level was calculated after 6 hours and 54 hours post heat treatment. Interestingly, even after 54 hours of treatment few var genes show slight upregulation in the expression level in comparison to control.

Other members of multivariant gene family like rifins and stevors are also known to play important role during the malaria pathogenesis. However, nothing much is known about their expression and role under different conditions. We studied expression of rifins and stevors under temperature stress condition. Interestingly, we observed a marked increase in number of rifins and stevors expressed during temperature stress conditions (Figure 6 A). Thus, in future studies temperature stress condition can be used to activate the expression of rifins and stevors to further characterize their role in *P. falciparum*.

## Discussions

Parasites constantly face different environment fluctuations during their life cycle [40]. These fluctuations can have different effect on the parasite growth and development. Most organisms have well developed stress response machinery which helps the organism to cope up the unfavourable conditions [41, 42]. Such machinery is not well studied and characterized in *Plasmodium falciparum*. Hence, cell-to-cell heterogeneity present in the population and how it affects the response of the parasites to different stress conditions is still unclear [8]. With recent studies appreciating the role of stress responses in drug resistance generation in *P. falciparum*, it is important to look at the effect of stress conditions on cell-to-cell variability using single cell RNA sequencing. Unlike previous scRNA-seq studies that were performed on mixed parasite populations [2, 28, 43] at different stages of growth, we performed scRNA sequencing of more number of cells at higher depth to better understand the composition and complexity of the synchronized *Plasmodium* cultures. Since synchronization is performed on a regular basis for different *in vitro* experiments, it is crucial to understand the transcriptional variation between sub-populations which can bias experimental outcomes. We found nine transcriptionally distinct cell populations all of which were represented in the Malaria Cell Atlas [28] (Supplementary Figure 1C). Thus, though parasites are morphologically synchronized, they exhibit a transcriptionally heterogeneous population.

Under temperature stress, few parasites (<1%) adopt a reactive state. They have higher expression level of stress responsive genes, gametocyte markers and multivariant gene family like var, rifins and stevor. Since rifins and stevor genes are expressed only in very few cells, not much is known about their role in parasitology. Therefore, temperature stress may be used to dissect the physiological and molecular role of rifins and stevors in *P. falciparum*. About 10% of the parasites are primed for stress adaptation under normal conditions. Stress responsiveness is an important determinant for drug resistance in *P. falciparum* [14, 15]. This is also evident from the fact that even a clonal population of resistant parasites do not show complete resistance (varying Ring Survival Assay (RSA) levels, a measure of resistance against artemisinin [44]. Thus, this sub-population (∼10% of the total population), which exhibits upregulation of stress response pathways might be responsible for the possible artemisinin resistance in the parasites. This finding was averaged out in previously bulk transcriptomic studies and may have hampered dissection of drug-resistance mechanism in *P. falciparum*.

Exposure to stress conditions can trigger the process of gametogenesis in *Plasmodium*. Here, we have identified a sup-population (< 5%) of parasites which express various gametocyte marker and regulator genes and thus committed for gametogenesis. Interestingly, this sub-population does not change upon temperature exposure but upregulates expression of gametocyte marker and regulator genes. This in turn suggests that a sub-population of the parasites is primed for gametogenesis and the temperature acts as cue to activate the marker genes and induces the gametogenesis. Furthermore, bulk RNA sequencing performed during temperature stress does not show any change in the levels of gametogenesis related genes indicating that there are very few parasites which are primed and identification of this rare population can only be done using single cell RNA sequencing techniques. Interestingly, we found a bifurcation for cell-fate decision to gametogenesis at an early and at a later stage of intra-erythrocytic cycle, illustrating that a sub-population of the parasites is always committed to gametogenesis. This may be a mechanism evolved by parasites to ensure continues transmission during each intra-erythrocytic cycle.

Thus, the scRNA-seq transcriptional profiling generated under temperature stress condition improved our understanding of the parasite stress response pathways under unfavourable condition. Our study also identifies the heterogeneity in the parasite population and how such diversity in their cellular states responds discreetly upon stress conditions. Importantly, the knowledge generated in this study could potentially be used to better understand the molecular mechanism of drug resistance, pathogenesis and virulence of the parasite.

## Materials and Methods

### *In Vitro* culture of *P. falciparum* parasites and temperature stress

*P. falciparum* strain 3D7 were cultured in RPMI1640 medium supplemented with 25 mM HEPES, 0.5 % AlbuMAX I, 1.77 mM sodium bicarbonate, 100 μM hypoxanthine and 12.5 μg ml-1 gentamicin sulfate at 37 °C. Parasites were maintained at 1-1.5% haematocrit and 5% parasitemia. Fresh O+ve human RBC’s were isolated from healthy human donor. Parasites were diluted after every two days by splitting the flask into 2-3 flasks in order to maintain the parasitemia around 5%. Parasites were synchronized at schizont stage using 63% percoll density gradient and synchronized at early ring stage using 5% sorbitol. Parasite growth was monitored using Giemsa staining of thin blood smear.

### Stress induction

Tightly synchronized parasites were given temperature stress for 6 hours from late ring stage (∼17 hours) to early trophozoite (∼23 hours) at 40° C. Control parasites were kept at normal culture conditions (37° C).

### Percoll gradient and isolation of single iRBCs

In order to get rid of the uninfected parasites we performed percoll gradient to separate uninfected RBC from the infected RBC. Different percentages of percoll gradient were tried separation of early trophozoite from uninfected since 63% percoll is used for separation of schizont from the uninfected parasites. 81% Percoll was finally used for the enrichment of the infected parasites after the experiment. Parasites were washed with 1X PBS twice to get rid of any dead parasites and parasites were counted before proceeding with bar-coding of the cells.

### scRNA-sequencing library preparation

Gel Beads-in-emulsion (GEMs) are generated by combining barcoded Single Cell 3’ v3 Gel Beads, a master mix containing cells and partitioning oil onto Chromium Chip B. Cells are delivered at a limiting dilution in order to attain single cell resolution. Once the gel beads is dissolved and cell are lysed inside the oil droplet, primers are released which binds to the poly A tail of the mRNAs and helps in the cDNA synthesis. Primers contain an Illumina TruSeq Read 1, 16 nucleotide 10x barcode, 12 nucleotide unique molecular identifier (UMI) and 30 nucleotide poly (dT) sequence. Enzymatic fragmentation and size selection are used to optimize the cDNA amplicon.

### scRNA-sequencing at NextSeq550 (Illumina)

Quality of the cDNA was estimated using Bioanalyzer and concentration of the sample was calculated using HS DNA Qubit. Transcriptome sequencing was performed using Illumina NextSeq 550 system (2x 75 bp read length) at Indian Institute of Science Education and Research, Pune.

### RNA sequencing and Data analysis

Bulk RNA sequencing was performed using parasites harvested for RNA isolation after the stress induction. Total RNA isolation was performed using TRIzol reagent. Bioanalyzer was performed to analyse the quality of RNA before proceeding for library preparation. Three biological replicates were pooled together for performing RNA sequencing. The cDNA libraries were prepared for samples using Agilent SureSelect strand specific RNA library preparation kit. Transcriptome sequencing was performed using Illumina NextSeq 550 system (2×150 bp read length) at Indian Institute of Science Education and Research, Pune. Quality control of the RNA-sequencing reads is performed using FASTQC and reads were trimmed based on the quality estimates. RNA paired directional reads were mapped to GTF annotation file v41.0 of the *P. falciparum* genome using TopHat. SAM-tools [45] were used for file handling and conversion. Cufflinks [46] (cuffmerge, cuffmerge and cuffdiff) programs were used for differential gene expression. MA plot is generated using ‘R’ software (http://r-project.org/). Gene ontology was performed using Plasmodb [47] (https://plasmodb.org/).

### scRNA-sequencing library preparation and sequencing

1000 cells/μL dilution of 80% viable cells was loaded onto the 10x chip for library preparation using Single cell v3 chemistry. The libraries for both the control and the treated sample were multiplexed together and were sequenced using mid output 75 bp single end configuration flow cell on the Illumina NextSeq 550 system. Cell ranger count was run on samples using v41.0 *P. falciparum* genome (PF3D7) to align and summarize per cell barcode (cell) read counts for each sample. For control samples 4949 cells were sequenced at a depth of 88.8 million reads with 88.1% reads in cells assigning a median of 17.9k reads/cell. For treated sample, 6873 cells were sequenced at a depth of 89.3 million reads with 88% reads in cells assigning a median of 13k reads/cell.

### scRNA-sequencing quality control and data analysis

The sparse matrices generated by 10x cell ranger count were read into R 3.5.3 using the Seurat 3.0.0 package. In order to remove potential duplets at a frequency of 5% and droplets with background RNA content, we removed cells with the top 5% and bottom 5% of total RNA molecules (or UMIs) from each sample. Thereafter, both the samples were combined using the Canonical correlation analysis using 10 principal components for finding integration anchors followed by 20 principal components to integrate the data. 20 principal components were used to perform t-SNE projections and clustering at a resolution of 0.5 using the Find Neighbours and Find Clusters functions (Supplementary Figure 3 and 4). Cluster specific markers were identified using the Find Conserved Markers function (with default parameters) and differentially expressed genes were identified using Find Markers function (with default parameters). Psuedotime analysis was performed using Monocle 2 package assuming a negbionomial distribution with a detection limit of 0.5 and a minimum normalized gene expression of 0.1. The marker genes for different clusters obtained from Seurat analysis above were used to calculate psuedotime and order the cells along psuedotime trajectory. Interactive web application and visualization was created using the open package from Rstudio called “Shiny apps” (https://www.shinyapps.io/).

### Patient sample data availability and analysis

Patient sample data analyzed from the microarray data obtained from NCBI’s Gene Expression Omnibus (GEO) with accession number GSE59099. Graphpad (www.graphpad.com) was used for plotting the normalized log2 expression ratios of the RNA sample/3D7 RNA reference pool for all probes. Single cell data was obtained from Malaria Cell Atlas (https://www.sanger.ac.uk/science/tools/mca).

### Primer design and RT-qPCR analysis

Total RNA was isolated from the parasites using TRIzol reagent (Biorad). Post DNAse treatment 2 μg RNA was used for cDNA synthesis using ImProm-II Reverse transcription system (Promega), as per the manufacture’s recommendation. Real time PCR was carried out using CFX96 Real Time PCR detection system (Biorad). 18S rRNA and tRNA synthetase were used as an internal control to normalize for variability across different samples. Quantification of the expression was done with the help of fluorescence readout of SYBR green dye incorporation into the amplifying targets (Biorad). Each experiment included technical triplicates and was performed over three independent biological replicates. Primers used for qPCR are mentioned in the Supplementary table S2.

### Var gene expression level

Violin plots (Figure 6C) was plotted for the number of cells expression different number of var genes. Maximum cells express single var genes per cell. But few cells also express higher number of var genes per cell. Number of cells expressing the one, two, three etc var genes were calculated and normalised with total number of cells in both control and temperature treatment. The normalized value was multiplied with 1000 and bar plots (Figure 6D) was plotted to indicate the different in the number of cells expression different number of var genes in control and temperature treatment.

## Data availability/submission

Single-cell RNA sequencing data for Plasmodium for control and temperature stress are submitted to Sequence Read Archive (SRA) under ID PRJNA560557.

## Ethics Statement

This study does not involve human participants. Human RBCs used in this study were obtained from the KEM Blood Bank (Pune, India) as blood from anonymized donors. Approval to use this material for *P. falciparum in vitro* culture has been granted by the Institutional Biosafety Committee of Indian Institute of Science Education and Research Pune (BT/BS/17/582/2014-PID).

## Conflict of interest

The authors declare that they have no conflict of interest.

## Acknowledgements

This work was supported by grants under DST INSPIRE (IFA-13, LMBM-53) and DBT-IYBA (BT/08/IYBA/2014-17) from Government of India to KK. MR is supported by DBT-SRF fellowship. We are thankful to Dr. Chandramouli Reddy for helpful discussions and NGS facility at IISER Pune for sequencing. The funders had no role in study design, data collection and analysis, decision to publish, or preparation of the manuscript.

## Supplementary Information

**Supporting table S1: Markers genes which are expressed at a higher level in each cluster. An excel sheet provided as additional information.**

**Supporting table S2:**
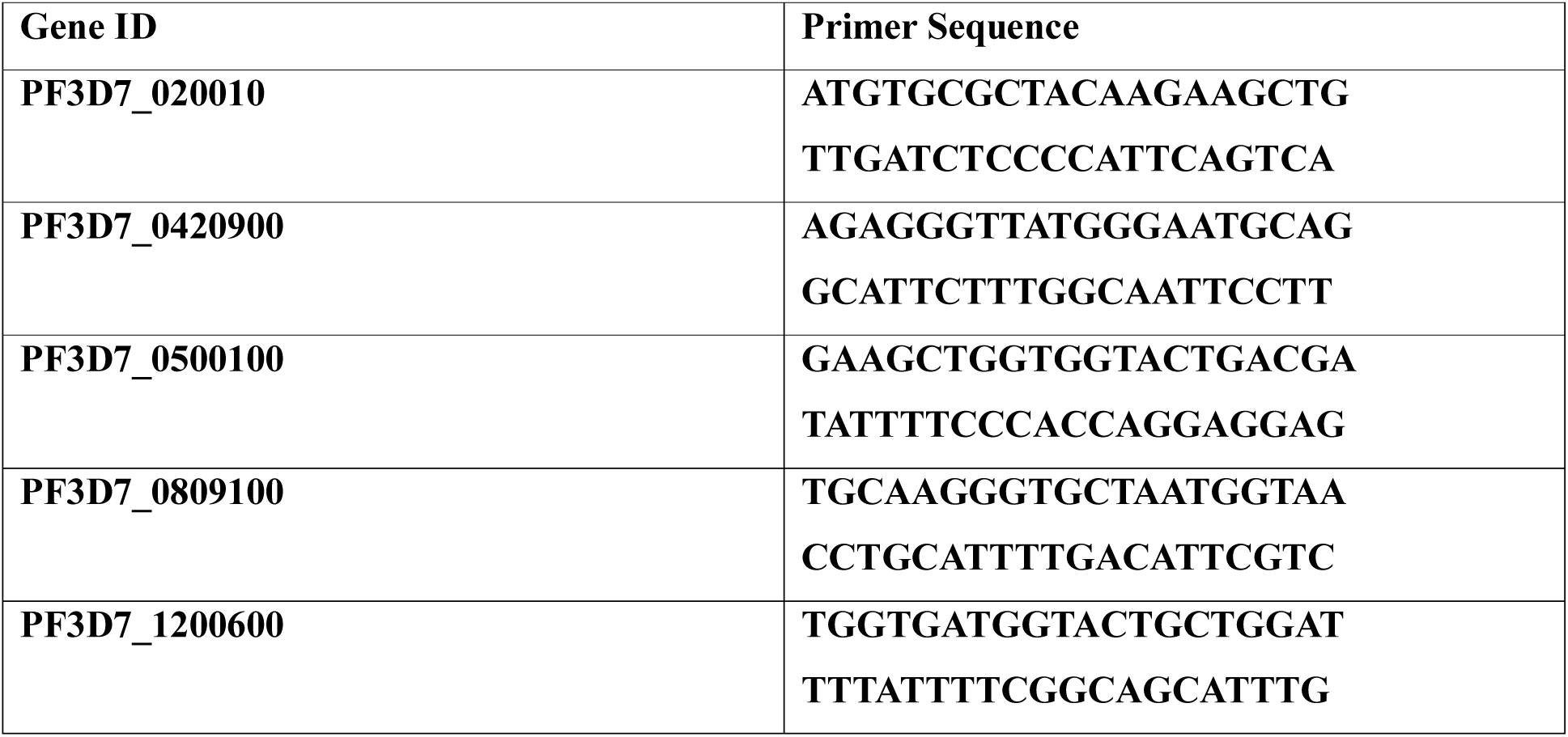
Primers used for RT-qPCR.

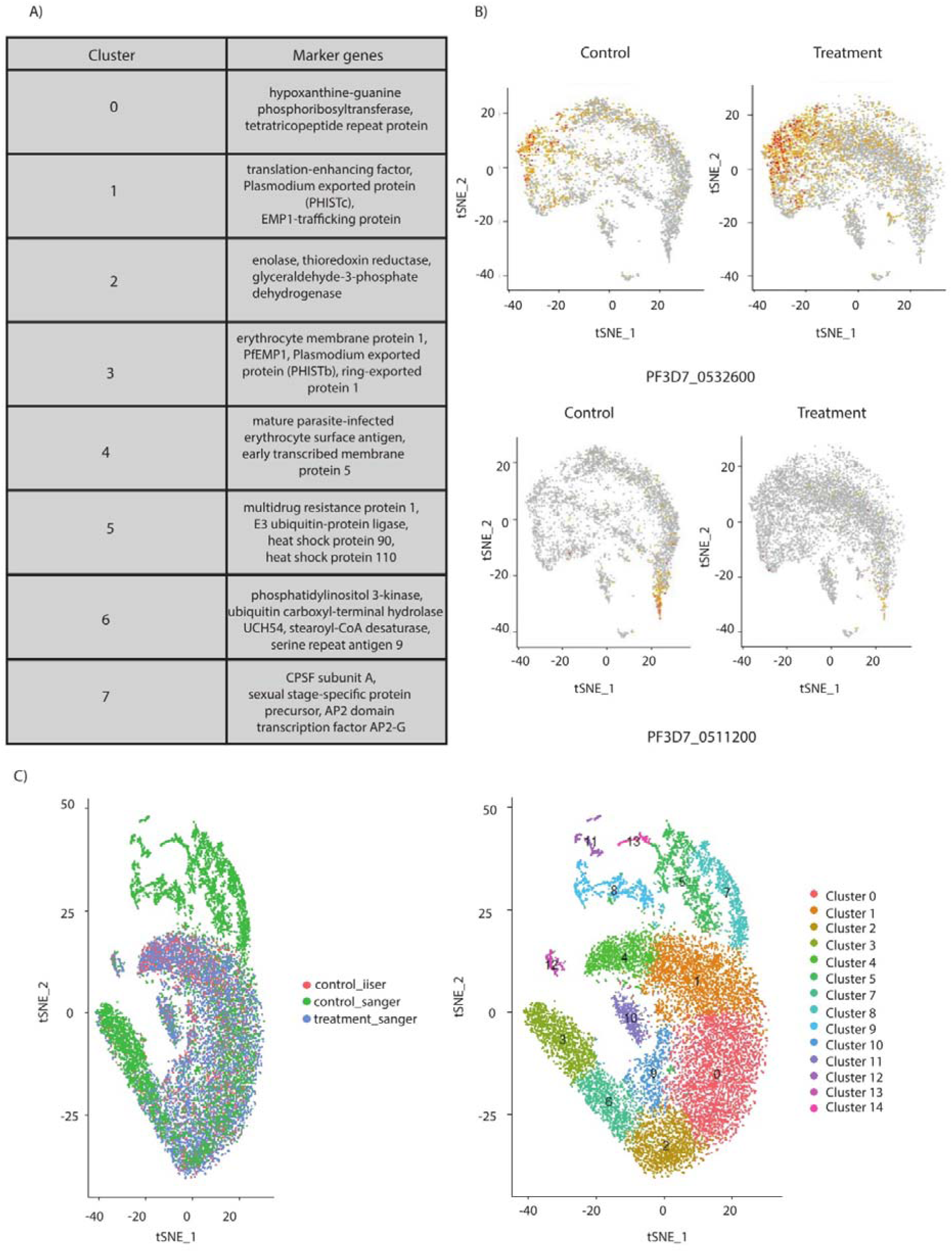

**Supplementary Figure 3:** A) Table showing the markers genes for each cluster obtained through single cell sequencing at control and temperature treatment. B) Representative tSNE plots showing the expression of PF3D7_0532600 and PF3D7_0511200 in control and treatment condition. C) Comparison of the clustering of control and treated sample with the control cells of data obtained from Howick et. al. using 10x Genomics.

**Supplementary Figure 2:**
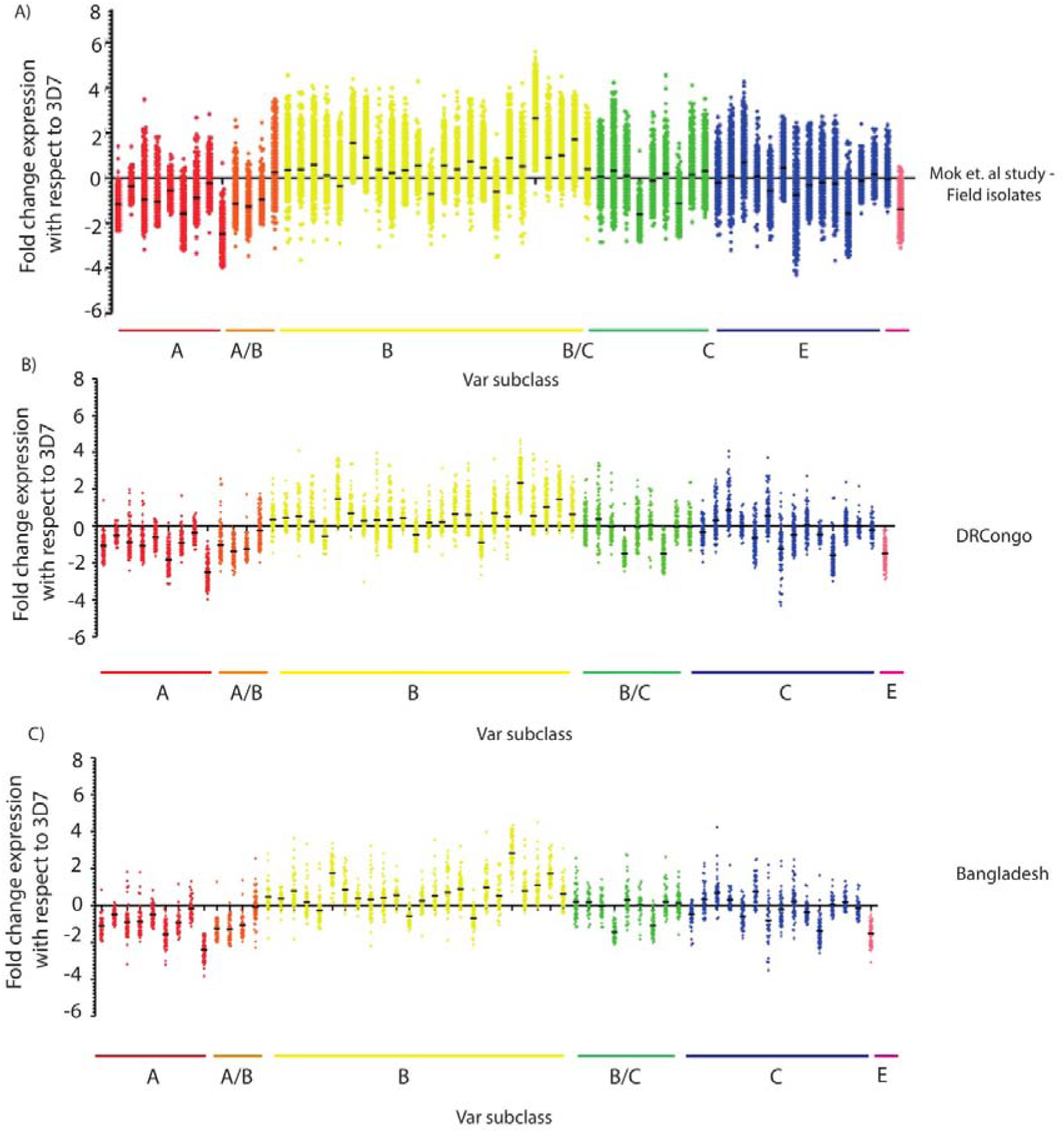
A) Fold change in expression level of the var genes in patient samples normalised to 3D7 strain. B) Fold change in expression level of the var genes in patients from Congo normalised to 3D7 strain. C) Fold change in expression level of the var genes in patients from Bangladesh normalised to 3D7 stain.

**Supplementary Figure S3:**
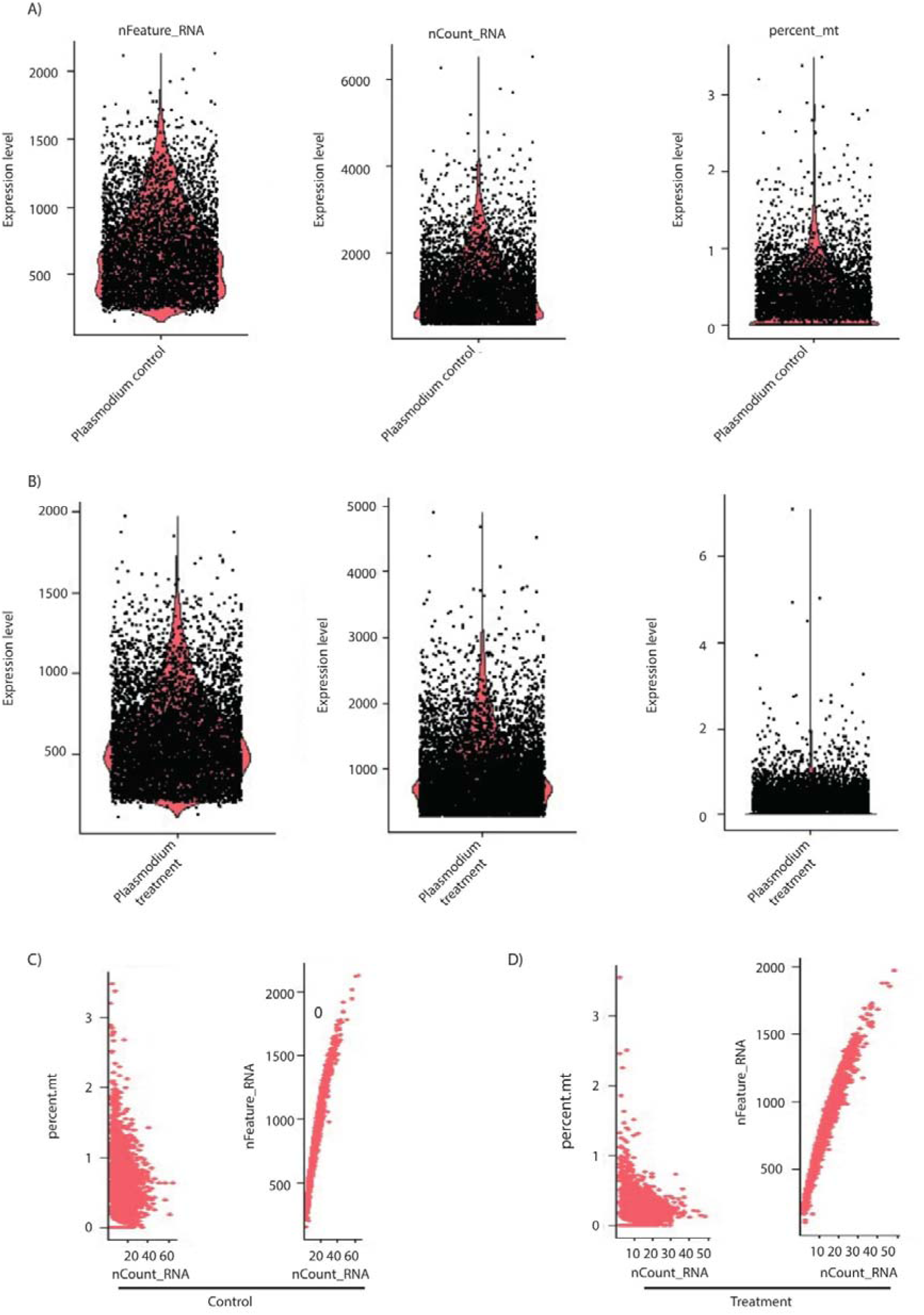
(A and B) Sanity check plots of number genes (nFeature_RNA), num ber of RNA molecules (nCount_RNA) and percentage of mitochondrial genes (genes beginning with “mal” in the annotation file) across individual cells in the dataset for both control and temperature stress sample respectively. (C and D) Sanity check plots to see any deviations from the expected relationship between number of RNA molecules (nCount_RNA) and percentage of mitochondrial gene expression (percent.mt); expected to be not correlated as shown; and number of genes (nFeature_RNA); expected to be linear because the deeper a cell is sequenced the more number of genes should be detected per cell.

**Supplementary Figure S4:**
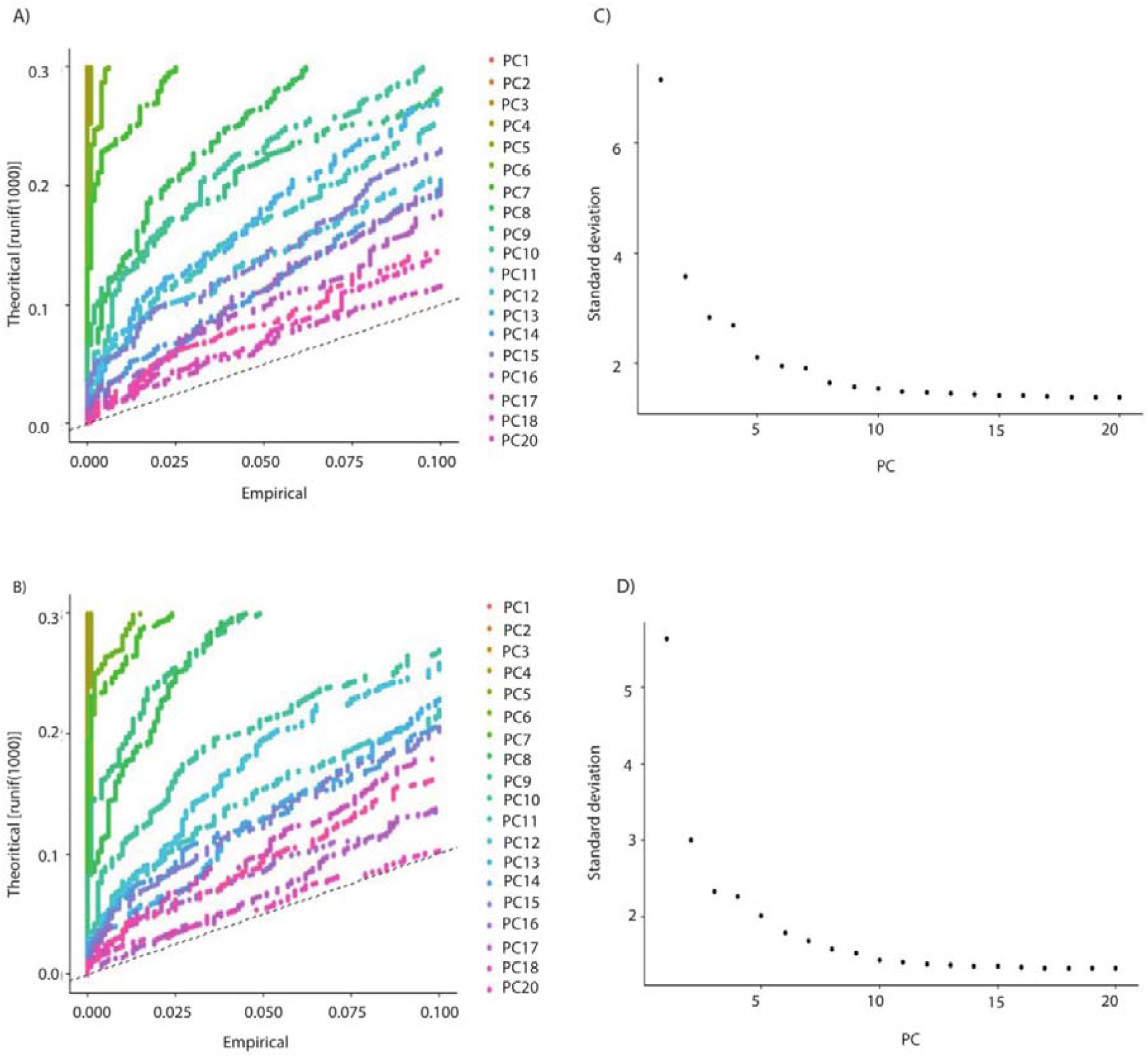
(A and B) JackStraw procedure was implemented to determine the dimensionality of the datasets (Macosko et al.). 10 significant principal components (PCs) with strong enrichment and low P-value were identified. (C and D) Elbow plots 6 (ranking of principal components based on the percentage of variance) of 20 principal components (PC) for the control and the treated sample show that the percentage variation explained per PC flattened around PC10. Hence, only 10 PCs were taken for downstream analysis like cell clustering. t-Distributed Stochastic Neighbor Embedding (t-SNE) projections of gene expression for control as well as treatment data set using nearest neighbourgraph based clustering followed by Louvain algorithm suggested 8 clusters across both data set. Both the datasets were combined to explore clusters which are common between the two conditions.

## References

1. WHO (2018) World Malaria Report.

2. Poran, A., Notzel, C., Aly, O., Mencia-Trinchant, N., Harris, C. T., Guzman, M. L., Hassane, D. C., Elemento, O. & Kafsack, B. F. C. (2017) Single-cell RNA sequencing reveals a signature of sexual commitment in malaria parasites, Nature. 551, 95–99.

3. Ngara, M., Palmkvist, M., Sagasser, S., Hjelmqvist, D., Bjorklund, A. K., Wahlgren, M., Ankarklev, J. & Sandberg, R. (2018) Exploring parasite heterogeneity using single-cell RNA-seq reveals a gene signature among sexual stage Plasmodium falciparum parasites, Experimental cell research. 371, 130–138.

4. Reid, A. J., Talman, A. M., Bennett, H. M., Gomes, A. R., Sanders, M. J., Illingworth, C. J., Billker, O., Berriman, M. & Lawniczak, M. K. (2018) Single-cell RNA-seq reveals hidden transcriptional variation in malaria parasites, Elife. 7, e33105.

5. Thien, H. V., Kager, P. A. & Sauerwein, H. P. (2006) Hypoglycemia in falciparum malaria: is fasting an unrecognized and insufficiently emphasized risk factor?, Trends Parasitol. 22, 410–5.

6. Pavithra, S. R., Banumathy, G., Joy, O., Singh, V. & Tatu, U. (2004) Recurrent fever promotes Plasmodium falciparum development in human erythrocytes, The Journal of biological chemistry. 279, 46692–9.

7. Percario, S., Moreira, D. R., Gomes, B. A., Ferreira, M. E., Goncalves, A. C., Laurindo, P. S., Vilhena, T. C., Dolabela, M. F. & Green, M. D. (2012) Oxidative stress in malaria, International journal of molecular sciences. 13, 16346–72.

8. Patra, P. & Klumpp, S. (2014) Phenotypically heterogeneous populations in spatially heterogeneous environments, Physical review E, Statistical, nonlinear, and soft matter physics. 89, 030702.

9. Smith, S. & Grima, R. (2018) Single-cell variability in multicellular organisms, Nature communications. 9, 345.

10. Martins, B. M. & Locke, J. C. (2015) Microbial individuality: how single-cell heterogeneity enables population level strategies, Curr Opin Microbiol. 24, 104–12.

11. Kwiatkowski, D. (1989) Febrile temperatures can synchronize the growth of Plasmodium falciparum in vitro, Journal of Experimental medicine. 169, 357–361.

12. Oakley, M. S., Kumar, S., Anantharaman, V., Zheng, H., Mahajan, B., Haynes, J. D., Moch, J. K., Fairhurst, R., McCutchan, T. F. & Aravind, L. (2007) Molecular factors and biochemical pathways induced by febrile temperature in intraerythrocytic Plasmodium falciparum parasites, Infect Immun. 75, 2012–25.

13. Udomsangpetch, R., Pipitaporn, B., Silamut, K., Pinches, R., Kyes, S., Looareesuwan, S., Newbold, C. & White, N. J. (2002) Febrile temperatures induce cytoadherence of ring-stage Plasmodium falciparum-infected erythrocytes, Proceedings of the National Academy of Sciences. 99, 11825–11829.

14. Rocamora, F., Zhu, L., Liong, K. Y., Dondorp, A., Miotto, O., Mok, S. & Bozdech, Z. (2018) Oxidative stress and protein damage responses mediate artemisinin resistance in malaria parasites, PLoS Pathog. 14, e1006930.

15. Dogovski, C., Xie, S. C., Burgio, G., Bridgford, J., Mok, S., McCaw, J. M., Chotivanich, K., Kenny, S., Gnadig, N., Straimer, J., Bozdech, Z., Fidock, D. A., Simpson, J. A., Dondorp, A. M., Foote, S., Klonis, N. & Tilley, L. (2015) Targeting the cell stress response of Plasmodium falciparum to overcome artemisinin resistance, PLoS Biol. 13, e1002132.

16. Abel, S. & Le Roch, K. G. (2019) The role of epigenetics and chromatin structure in transcriptional regulation in malaria parasites, Brief Funct Genomics.

17. Batugedara, G., Lu, X. M., Bunnik, E. M. & Le Roch, K. G. (2017) The Role of Chromatin Structure in Gene Regulation of the Human Malaria Parasite, Trends Parasitol. 33, 364–377.

18. Karmodiya, K., Pradhan, S. J., Joshi, B., Jangid, R., Reddy, P. C. & Galande, S. (2015) A comprehensive epigenome map of Plasmodium falciparum reveals unique mechanisms of transcriptional regulation and identifies H3K36me2 as a global mark of gene suppression, Epigenetics & chromatin. 8, 32.

19. Gupta, A. P., Chin, W. H., Zhu, L., Mok, S., Luah, Y. H., Lim, E. H. & Bozdech, Z. (2013) Dynamic epigenetic regulation of gene expression during the life cycle of malaria parasite Plasmodium falciparum, PLoS Pathog. 9, e1003170.

20. Coleman, B. I. & Duraisingh, M. T. (2008) Transcriptional control and gene silencing in Plasmodium falciparum, Cell Microbiol. 10, 1935–46.

21. Cui, L., Lindner, S. & Miao, J. (2015) Translational regulation during stage transitions in malaria parasites, Annals of the New York Academy of Sciences. 1342, 1–9.

22. Vembar, S. S., Droll, D. & Scherf, A. (2016) Translational regulation in blood stages of the malaria parasite Plasmodium spp.: systems-wide studies pave the way, Wiley interdisciplinary reviews RNA. 7, 772–792.

23. Foth, B. J., Zhang, N., Chaal, B. K., Sze, S. K., Preiser, P. R. & Bozdech, Z. (2011) Quantitative time-course profiling of parasite and host cell proteins in the human malaria parasite Plasmodium falciparum, Molecular & Cellular Proteomics. 10, M110. 006411.

24. Choi, Y. H. & Kim, J. K. (2019) Dissecting Cellular Heterogeneity Using Single-Cell RNA Sequencing, Molecules and cells. 42, 189–199.

25. Peng, J., Sun, B. F., Chen, C. Y., Zhou, J. Y., Chen, Y. S., Chen, H., Liu, L., Huang, D., Jiang, J., Cui, G. S., Yang, Y., Wang, W., Guo, D., Dai, M., Guo, J., Zhang, T., Liao, Q., Liu, Y., Zhao, Y. L., Han, D. L., Zhao, Y., Yang, Y. G. & Wu, W. (2019) Single-cell RNA-seq highlights intratumoral heterogeneity and malignant progression in pancreatic ductal adenocarcinoma, Cell research.

26. Cheng, S., Pei, Y., He, L., Peng, G., Reinius, B., Tam, P. P. L., Jing, N. & Deng, Q. (2019) Single-Cell RNA-Seq Reveals Cellular Heterogeneity of Pluripotency Transition and X Chromosome Dynamics during Early Mouse Development, Cell reports. 26, 2593–2607 e3.

27. Conover, W. J., Johnson, M. E. & Johnson, M. M. (1981) A comparative study of tests for homogeneity of variances, with applications to the outer continental shelf bidding data, Technometrics. 23, 351–361.

28. Howick, V. M., Russell, A., Andrews, T., Heaton, H., Reid, A. J., Natarajan, K. N., Butungi, H., Metcalf, T., Verzier, L. H. & Rayner, J. (2019) The Malaria Cell Atlas: a comprehensive reference of single parasite transcriptomes across the complete Plasmodium life cycle, BioRxiv, 527556.

29. Carter, R. & Miller, L. H. (1979) Evidence for environmental modulation of gametocytogenesis in Plasmodium falciparum in continuous culture, Bulletin of the World Health Organization. 57, 37–52.

30. Dyer, M. & Day, K. (2000) Commitment to gametocytogenesis in Plasmodium falciparum, Parasitology today. 16, 102–107.

31. Smalley, M. & Brown, J. (1981) Plasmodium falciparum gametocytogenesis stimulated by lymphocytes and serum from infected Gambian children, Transactions of the Royal Society of Tropical Medicine and Hygiene. 75, 316–317.

32. Buckling, A., Ranford-Cartwright, L., Miles, A. & Read, A. F. (1999) Chloroquine increases Plasmodium falciparum gametocytogenesis in vitro, Parasitology. 118, 339–346.

33. Trager, W. & Gill, G. S. (1992) Enhanced gametocyte formation in young erythrocytes by Plasmodium falciparum in vitro, The Journal of protozoology. 39, 429–432.

34. Sokhna, C., Trape, J. & Robert, V. Gametocytaemia in Senegalese children with uncomplicated falciparum malaria treated wit h chloroquine, amodiaquine or sulfadoxine plus pyrimethamine, Parasite I. 200, 8.

35. Stephens, J. W. W. & Christophers, S. R. (1908) The practical study of malaria and other blood parasites, University Press of Liverpool.

36. Kyes, S. A., Kraemer, S. M. & Smith, J. D. (2007) Antigenic variation in Plasmodium falciparum: gene organization and regulation of the var multigene family, Eukaryotic cell. 6, 1511–1520.

37. Rosenberg, E., Ben-Shmuel, A., Shalev, O., Sinay, R., Cowman, A. & Pollack, Y. (2009) Differential, positional-dependent transcriptional response of antigenic variation (var) genes to biological stress in Plasmodium falciparum, PLoS One. 4, e6991.

38. Hakimi, M.-A. & Deitsch, K. W. (2007) Epigenetics in Apicomplexa: control of gene expression during cell cycle progression, differentiation and antigenic variation, Current opinion in microbiology. 10, 357–362.

39. Ubhe, S., Rawat, M., Verma, S., Anamika, K. & Karmodiya, K. (2017) Genome-wide identification of novel intergenic enhancer-like elements: implications in the regulation of transcription in Plasmodium falciparum, BMC genomics. 18, 656.

40. Arnoldini, M., Mostowy, R., Bonhoeffer, S. & Ackermann, M. (2012) Evolution of stress response in the face of unreliable environmental signals, PLoS computational biology. 8, e1002627.

41. Dhar, R., Sägesser, R., Weikert, C. & Wagner, A. (2012) Yeast adapts to a changing stressful environment by evolving cross-protection and anticipatory gene regulation, Molecular biology and evolution. 30, 573–588.

42. Young, J. W., Locke, J. C. & Elowitz, M. B. (2013) Rate of environmental change determines stress response specificity, Proceedings of the National Academy of Sciences. 110, 4140–4145.

43. Walzer, K. A., Kubicki, D. M., Tang, X. & Chi, J.-T. A. (2018) Single-cell analysis reveals distinct gene expression and heterogeneity in male and female Plasmodium falciparum gametocytes, mSphere. 3, e00130–18.

44. Mbengue, A., Bhattacharjee, S., Pandharkar, T., Liu, H., Estiu, G., Stahelin, R. V., Rizk, S. S., Njimoh, D. L., Ryan, Y. & Chotivanich, K. (2015) A molecular mechanism of artemisinin resistance in Plasmodium falciparum malaria, Nature. 520, 683.

45. Li, H., Handsaker, B., Wysoker, A., Fennell, T., Ruan, J., Homer, N., Marth, G., Abecasis, G., Durbin, R. & Genome Project Data Processing, S. (2009) The Sequence Alignment/Map format and SAMtools, Bioinformatics. 25, 2078–9.

46. Trapnell, C., Roberts, A., Goff, L., Pertea, G., Kim, D., Kelley, D. R., Pimentel, H., Salzberg, S. L., Rinn, J. L. & Pachter, L. (2012) Differential gene and transcript expression analysis of RNA-seq experiments with TopHat and Cufflinks, Nature protocols. 7, 562–78.

47. Bahl, A., Brunk, B., Crabtree, J., Fraunholz, M. J., Gajria, B., Grant, G. R., Ginsburg, H., Gupta, D., Kissinger, J. C., Labo, P., Li, L., Mailman, M. D., Milgram, A. J., Pearson, D. S., Roos, D. S., Schug, J., Stoeckert, C. J., Jr. & Whetzel, P. (2003) PlasmoDB: the Plasmodium genome resource. A database integrating experimental and computational data, Nucleic acids research. 31, 212–5.

